# Stochastic dynamics of ecological populations subjected to environmental perturbations

**DOI:** 10.1101/2023.02.16.528890

**Authors:** Sayeh Rezaee, Cesar Nieto, Zahra Vahdat, Abhyudai Singh

**Affiliations:** Department of Electrical and Computer Engineering, University of Delaware, Newark, DE, USA; Department of Electrical and Computer Engineering, Biomedical Engineering, Mathematical Sciences, Center of Bioinformatic and Computational Biology, University of Delaware, Newark, DE, USA

## Abstract

The stochastic logistic model is widely used to capture random fluctuations arising from birth-death processes in ecological populations. We use this model to study the impact of environmental perturbations that may occur naturally or as a consequence of population harvesting. In our model formulation, environmental perturbations occur randomly as per a Poisson process, and the perturbations result in each individual dying with a certain probability of death. Moment closure schemes are employed to derive expressions for the mean and variability in population numbers. Moreover, to quantify the impact of population extinction in our model we compute the probability of extinction using the Finite State Projection (FSP) numerical method. Our analysis shows that rare environmental perturbations with a high probability of death lead to overall larger random fluctuations and extinction risk as compared to frequent perturbations with a low probability of death. Finally, we formulate the problem in the context of population harvesting to find the optimal harvesting rate that maximizes the cumulative yield.

## I. Introduction

There is a rich literature on modeling fluctuations in ecological population dynamics incorporating the effects of both environmental and demographic stochasticity [1]–[6]. Perhaps the most useful framework for doing it is the stochastic logistic model where a population *x* grows exponentially with a rate *r*, and the limitation of resources is captured by a death rate that scales linearly with population size. In the deterministic limit, the time evolution of the population density *x*(*t*) is given by the ordinary differential equation (ODE):

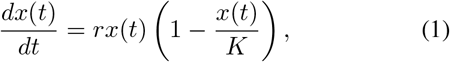

and asymptotically approaches the carrying capacity *K* [7]. The stochastic counterpart of this process is based on probabilistic birth/death events that increase/decrease the integer-valued population count by one, respectively. In contrast to the deterministic approach, the population will eventually always go extinct (i.e. *x* = 0 with probability one) in the stochastic approach, although for many parameter regimes the time to extinction may be large [8].

Our goal here is to use this framework to investigate the impact of environmental perturbations that occur via natural disasters or through harvesting of the population. Previous work has used the stochastic logistic model to describe the evolution of ecological populations under random environmental perturbations [9]–[12]. However, much of this has relied on a stochastic differential equation framework of additive noise. In this contribution, we consider the environmental events to occur as per a constant Poisson rate. Let us assume a specific probability of mortality for each individual. Whenever the event occurs, the surviving population is modeled as a Binomial distribution, conditioned on the size of population just before the event. In the limit of larger population sizes, where the mean time of extinction is large, we develop approximate analytical formulas quantifying the magnitude of fluctuations in population numbers. Then we apply finite state projection (FSP) method [13] to explicitly compute the probability of extinction. Comparing these two methods, it is evident that the proposed model gives a satisfactory approximation, given a biologically rational range for the parameters and time.

Next, we use this formulation to study the optimal harvesting problem, where a trade-off is set up based on overexploitation leading to population extinction. The optimal strategies for exploiting biological resources have been one of the significant topics of research since the amount and frequency of harvesting is important in both senses of biology and economy. There is a large literature on analyzing the harvesting problem [14]–[16]. Previous contributions [17], [18] have studied the optimal harvesting strategies for deterministic systems, which is reasonable for populations with relatively large carrying capacities where the probability of extinction can be neglected. Having this assumption, the harvesting problem can be modeled using a stationary distribution for the population [8]. However, given that environmental randomness can significantly affect the population dynamics, many studies were done on stochastic models for harvesting [19]–[21].

Our approach revolves around introducing a second variable we refer to as “yield’ that increases by a jump whenever the population is harvested based on a given constant Poisson rate. The yield also decreases exponentially (due to consumption) between successive harvesting events. We explore this two-dimensional stochastic system using the approach of moment dynamics [22]–[25]. However, model nonlinearities lead to an unclosed system (i.e., the time evolution of lower-order moments depends on the higher-order ones). Employing closure schemes that express high-order moments in terms of lower-order moments [26]–[33], we solve for average total yield analytically and find the optimal harvesting rate, at which the total yield is maximized. Then by comparing to numerical results from the FSP method, we discuss how population extinction events can alter analytical results obtained from moment dynamics.

## II. Stochastic logistic model with environmental perturbations

In this section, we will introduce the stochastic formulation of the population dynamics as a birth-death process with environmental perturbations as random jumps. We present different levels of analytical approximations of the solution for the population statistical moments. This, together with a comparison of the derived formulas with numerical methods which reach a more accurate solution, helps us to estimate the effect of these perturbations on the population variability.

Let the random process *x*(*t*) ∈ {0, 1, 2,…} denote the population count of a given species at time *t*. The process evolves as a result of three classes of probabilistic events.

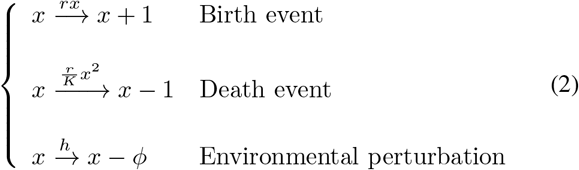

Each event occurs as per a rate given on the top of the arrow, and when the event happens, the population is reset as per the corresponding reset map. For example, the probability of a birth event in the next infinitesimal time interval (*t, t + dt*] is *rx*(*t*)*dt*, and this event increases the population number by one. The last event models the environmental perturbations occurring with a constant rate *h*, in which case *ϕ* individuals die (or harvested). The random variable *ϕ* is drawn from a Binomial distribution with parameters *x* and *f*, where each of *x* individuals die with probability *f*.

### A. Analytical Approximation

To study the population mean and variability in steady state, we present the system statistical moment dynamics and solve it in steady state. It should be noted that, in the stochastic model, the stationary distribution is degenerate at the origin with probability one, which is equivalent to the population extinction [30]. Yet, we can consider the stochastic model with non-extinct population by assuming a sufficiently large enough time to extinction. Considering this assumption, we approximately find the population statistics for the stochastic model. Due to being discrete, we expect this model to be closer to the reality, comparing to the continuous deterministic model. Later throughout the article, we solve the system numerically while considering the probability of extinction, and we discuss how good our analytical approximation works.

The time evolution of first- and second-order moments for the stochastic system (2), can be obtained by using the corresponding Dynkin’s formula [34], where 〈〉 denotes the expected value operation.

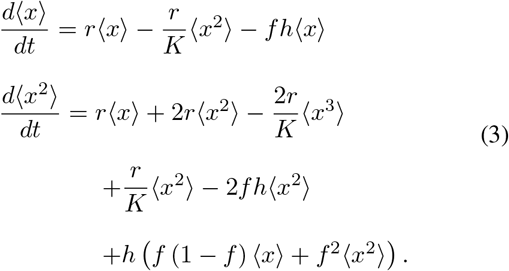

Before calculating the stochastic system’s moments, we can derive the deterministic approximation of the system’s solution. This deterministic limit consists of approximating *x* as a deterministic variable, which is equivalent to taking 〈*x*^2^〉 ≈ 〈*x*〉^2^, and leads to the following ODE.

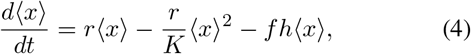

From the above equation, the steady-state population level of the deterministic approximation, denoted by *K_e_*, is given.

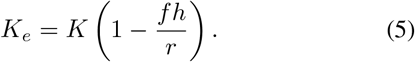

Now, we proceed by solving the moments obtained from stochastic formulation. Note the appearance of high-order moments on the right-hand-side of (3). This leads to a system of unclosed moment dynamics arising due the nonlinear propensity of the death event. Previously, we had developed a derivative-matching moment closure scheme for the stochastic logistic model [35], which approximates

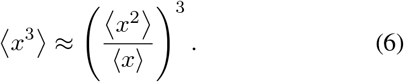

Using this approximation in (3), we obtain the following first- and second-order moments at steady state.

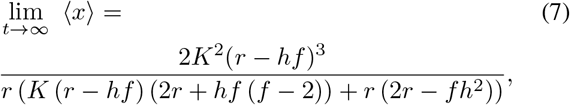

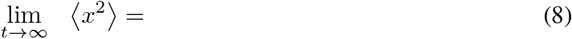

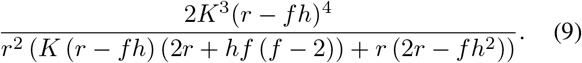

As we are interested in investigating how the frequency of perturbation events *h* and the probability of death *f* affect the population, we quantify the population variability by steadystate coefficient of variation *CV*^2^, which is described as

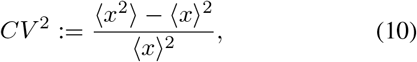

and yields the following equation.

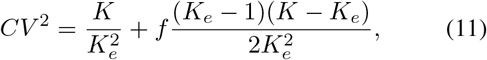

where *K_e_* is the steady-state population level in the deterministic approximation as predicted by the ODE in (5). In the absence of environmental perturbations, we have *f* = 0 and *K_e_* = *K*, which leads to

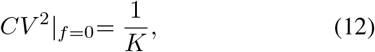

as previously reported [35], and it intuitively makes sense with the extent of fluctuations decreasing with increasing population size. However, in the presence of environmental perturbations (i.e. *f* ≠ 0), the fluctuation *CV*^2^ depends on both *h* and *f*. We plot the steady-state *CV*^2^ (11) as a function of increasing *f*, while the frequency of perturbation events *h* is decreasing such that *K_e_* remains fixed. As it is shown in Fig. 1.A, increasing the probability of death *f*, increases the *CV*^2^ for a fixed *K_e_*. Thus, increasing severity of events (while decreasing the frequency of events *h*) will drive larger fluctuations for the same average population size, which leads to higher risk of extinction. In the next section, we numerically solve the stochastic system to compare with the analytical result obtained in this section.

**Fig. 1:**
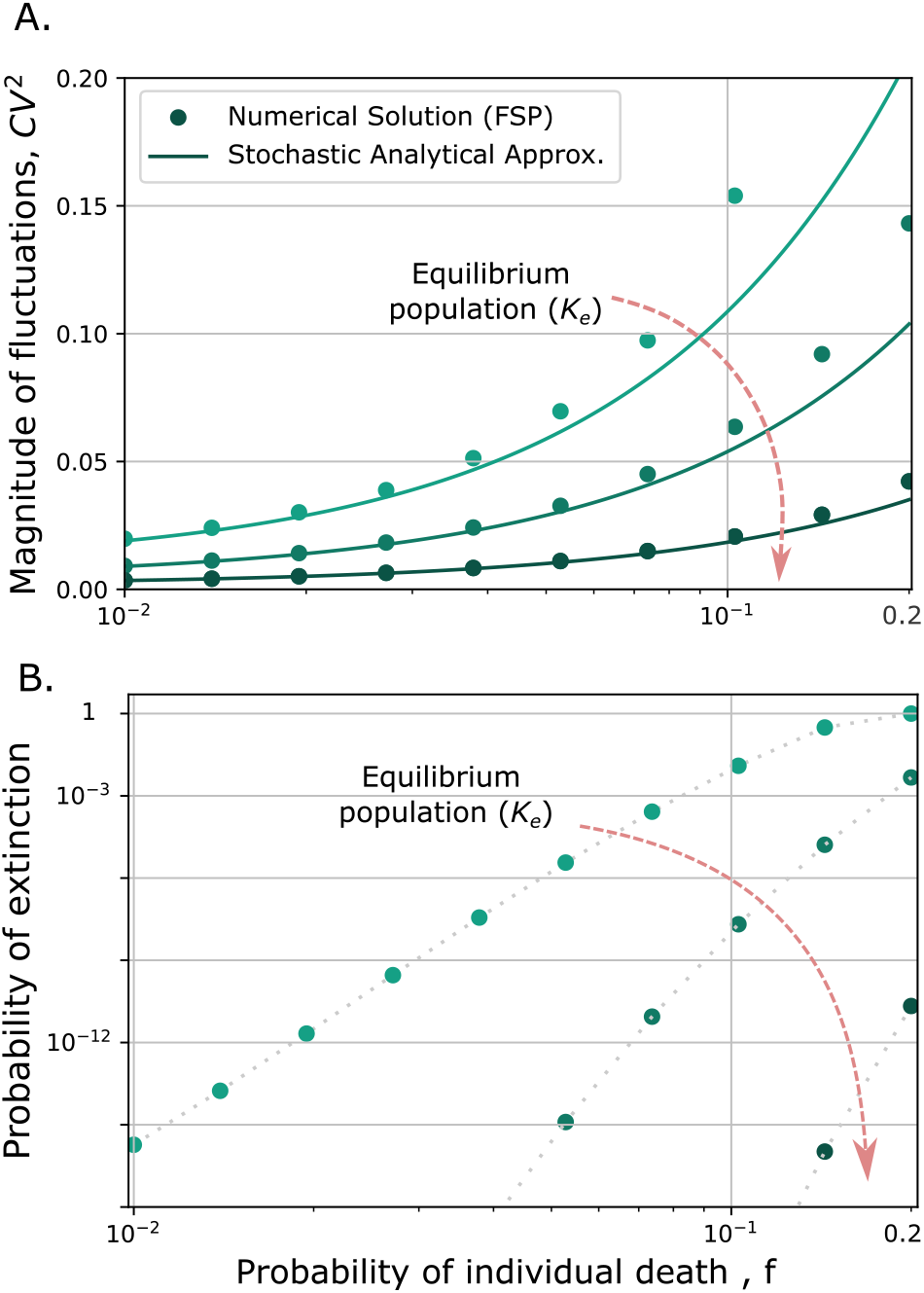
Rare environmental perturbations enhance fluctuations in population counts and the probability of extinction. A. Plot of the steadystate coefficient of variation (11) as a function of increasing *f*. The dots are achieved from the solution of FSP in (17) while *t_max_* = 400/*r*. B. Diagram of probability of extinction as a function of increasing *f* obtained from numerically solving (17) from FSP at a time *t_max_ =* 400/*r*. For both plots, the frequency of perturbation is decreased to keep *K_e_* fixed. Lines are made for different population counts as *K_e_* = *K*/3, *K*/2, 3*K*/4, increasing in arrow direction. Other parameters are taken as *r* = 10, *K* = 10^3^.

### B. Numerical Solution

The birth-death process described in (2) can be modeled using the forward Chapman-Kolmogorov formulation [36], also known as master equation. In this formalism, we define the probability vector *P*(*t*) with components *p_i_*(*t*) referring to the probability of having i individuals at time *t*. In general, 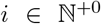 makes *P*(*t*) a vector with an infinite number of components. To achieve a numerical approximation of the master equation solution, we propose to use the finite state projection (FSP) method [13]. FSP is a numerical algorithm consisting on the truncation of the infinite number of equations in the master equation. Hence, if we consider only the first *I* elements of *P*(*t*) such that

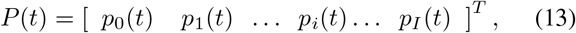

the master equation associated to the system gives

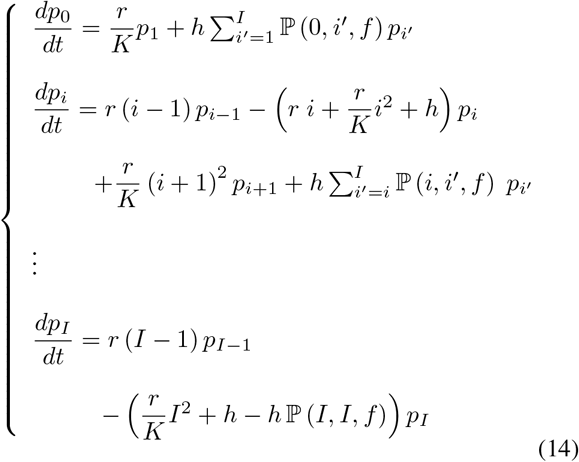

Note that the equations above utilize the probability mass function of the Binomial distribution denoted as 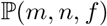.

This function represents the probability of obtaining a specific number of successes *m*, in a series of independent Bernoulli trials *n*, where the probability of success in each trial is denoted as *f*.

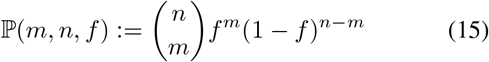

We aim to calculate the vector of probabilities defined in (13). To do so, we can express the dynamics of *P*(*t*) as a linear system in the form of

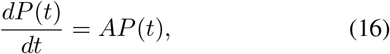

where *A*_(*I*+1)×(*I*+)_ is a square matrix derived from (14). Hence, *P*(*t*) is achieved by solving the above equation as

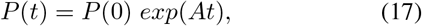

where *P*(0) = [0 0 … 1 … 0 0] is the vector of probabilities at time *t* = 0. Now, we numerically calculate *P*(*t*), in which *p*_0_(*t*) gives us the probability of extinction. In addition, the *n* – th order moment of *x* denoted by 〈*x^n^*〉 can be calculated from

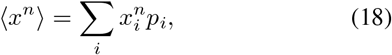

where *p_i_* is the probability of having *i* population count, and is obtained from (17). Accordingly, having the values for moments of *x, CV*^2^ can be achieved using its definition in (10). Fig. 1.A represents the magnitude of population fluctuations *CV*^2^ obtained numerically, along with the one computed from the analytical approximation using (11), as a function of the probability of death *f*. The result shows that for smaller values of stationary population level *K_e_* and greater values of *f*, the proposed analytical approximation is less accurate, as we assumed having a nonextinct population, while the probability of extinction in this range of parameters is relatively high (see Fig. 1.B). Additionally, this figure shows the probability of extinction achieved from the FSP solution, as a function of individual death probability *f*, while the frequency of perturbation *h* is decreased to keep *K_e_* fixed. As we have claimed earlier, the result in this figure shows that rare environmental perturbations with a higher probability of death increase the probability of extinction.

## III. Modeling harvesting with constant poisson rate

In the previous section, we investigated the stochastic logistic model with perturbations and the effect of frequency and severity of perturbation events on population variability. Now, we apply this model to consider environmental perturbations as harvesting events, which occur at a Poisson constant rate *h*. In the proposed model, each individual has a probability *f* of being harvested. In experiments, *f* can be estimated from the details of the harvesting method. Subsequently, our aim is to study the population statistics and, given *f*, find the harvesting rate that maximizes the average total yield.

### A. Deterministic Formulation of the Harvesting Model

Let us start by discussing the yield and its optimal value for the deterministic harvesting model. As was explained in the previous section, the stochastic logistic model with perturbations can be written as below in the deterministic limit.

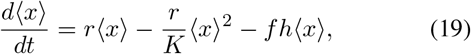

while here, we define *f* as the harvesting ratio, and *h* as the harvesting rate. In addition, we consider the dynamics of the total yield *y* to be continuously decreased by the consumption rate *λ*, and increased by the value of *fhx* due to harvest.

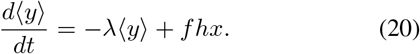

The steady-state total yield can be calculated from solving (19) and (20) in steady state as follows.

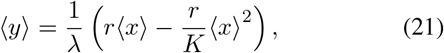

From the above equation, We can find the optimal harvesting rate *h*\* at which the total yield is maximized.

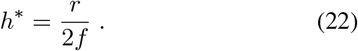

Hence, the corresponding maximum total yield is

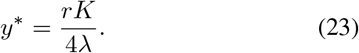

This value is the deterministic limit of the optimal total yield. However, continuous harvesting is an idealization and a more realistic modeling should include discrete jumps. Accordingly, we propose the stochastic harvesting model in the next section.

### B. Stochastic Formulation of the Harvesting Model

We develop the stochastic model explained in (2) such that the harvesting events are modeled by stochastic jumps occurring at a Poisson rate h. Every time harvesting occurs, each individual is caught with a probability *f*, such as the population *x* decreases by a quantity *ϕ*, which is a random variable derived from a Binomial distribution with parameters *x* and *f*, and each of *x* individuals die with probability *f*. The same value is added to the total yield *y* by each harvest. Hence, we can show the reset map of the model as follows.

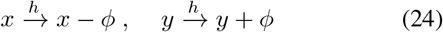

with 0 < *f* < 1 defining the harvesting ratio. Intuitively, in the case of having constant Poisson rate, the probability of harvest in an infinitesimal time is a constant. Additionally, the time between consecutive harvests follows an exponential distribution with mean 1/*h*.

Next, we assume that the total yield exponentially decreases by the consumption rate *λ*, leading to following dynamics.

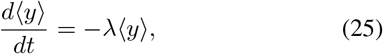

The schematic model of the described stochastic hybrid system is illustrated in Fig. (2).

**Fig. 2:**
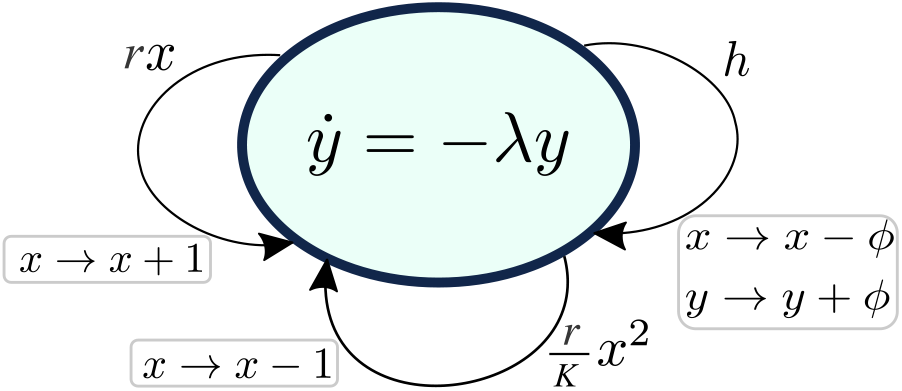
Schematic model of harvesting as a stochastic hybrid system. In this model, the changes in population count *x* are represented by stochastic jumps, and the yield *y* decreases exponentially over time due to a consumption rate of *λ*. Additionally, each harvest happens at a rate of *h* and results in a reduction of the population by a random value following a binomial distribution with a parameter of *ϕ*, while the yield increases by the same amount.

To study optimal harvesting, we define the optimal harvesting rate as the one that maximizes total yield. Hence, for our stochastic system, we need to find the maximum value of the mean yield 〈*y*〉. As explained in previous sections, to find the statistical moments of a random variable, we can write moment dynamics using Dynkin’s formula [34] and find the moments in steady state. Thus, from (24) and (25) we have

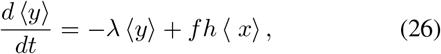

Consequently, the steady-state first-order moment of *y*, i.e., the steady-state mean yield is

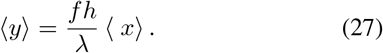

The goal here is to find the optimal harvesting rate and the corresponding maximum yield, which according to the above equation equals to

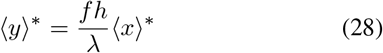

The steady-state first-order moment of *x*, indicated by 〈*x*〉, was already obtained in section II, where we studied the statistics of stochastic logistic model with perturbations. Since we are approximating the statistics at the limit of larger population sizes, which also leads to 〈*x*^2^〉 ≫ 〈*x*〉, we can use the below approximation to simplify moment dynamics in (3).

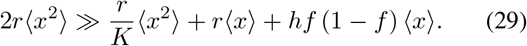

This approximation yields to the following steady-state first- and second-order moments of *x*.

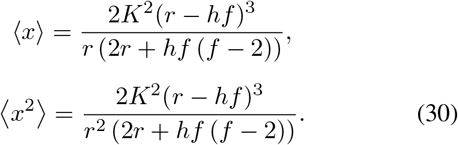

Consequently, we obtain the steady-state mean of total yield.

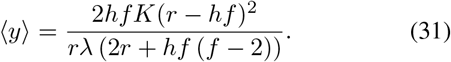

Now, we can calculate the optimal harvesting rate *h\** which maximizes the mean total yield.

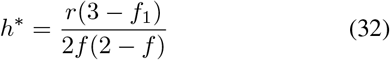

where 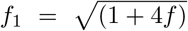. Additionally, the corresponding optimal yield 〈*y*〉* is

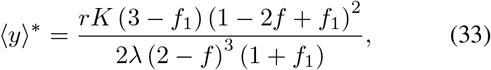

which is a decreasing function of *f*, so given 0 < *f* < 1, 〈*y*〉* is maximized when *f* = 0. This upper limit for the stochastic model is equal to the deterministic limit for 〈*y*〉^*^ which was obtained in (23). Additionally, it can be simply shown that the optimal harvesting rate obtained for the stochastic model in (32) is always smaller than the optimal harvesting rate achieved for deterministic model in (22). Consequently, incorporating stochasticity in the model resulted in a reduction of both the optimal harvesting rate and the maximum yield, making them more accurate comparing to the deterministic model (see Fig. 3). This figure shows the population statistics obtained in this section. To compare this approximation with the case in which extinction is not neglected, we use the results of the FSP method explained in Section II-B. The mean population 〈*x*〉 obtained numerically from this approach is represented in Fig. 3.B. As it is observable, the effects of extinction are important for higher values of *h* which makes the approximation less accurate for this parameter regime. However, noticing the optimal values of *k* and 〈*y*〉 in 3.C, we can see that extinction has a relatively less effect at the optimal harvest rate. In addition to the effects of the extinction, we can also observe how population variability can affect harvesting. The effect of stochasticity can be understood by comparison of the results of the continuous deterministic (green) and the stochastic approximation (blue).

**Fig. 3:**
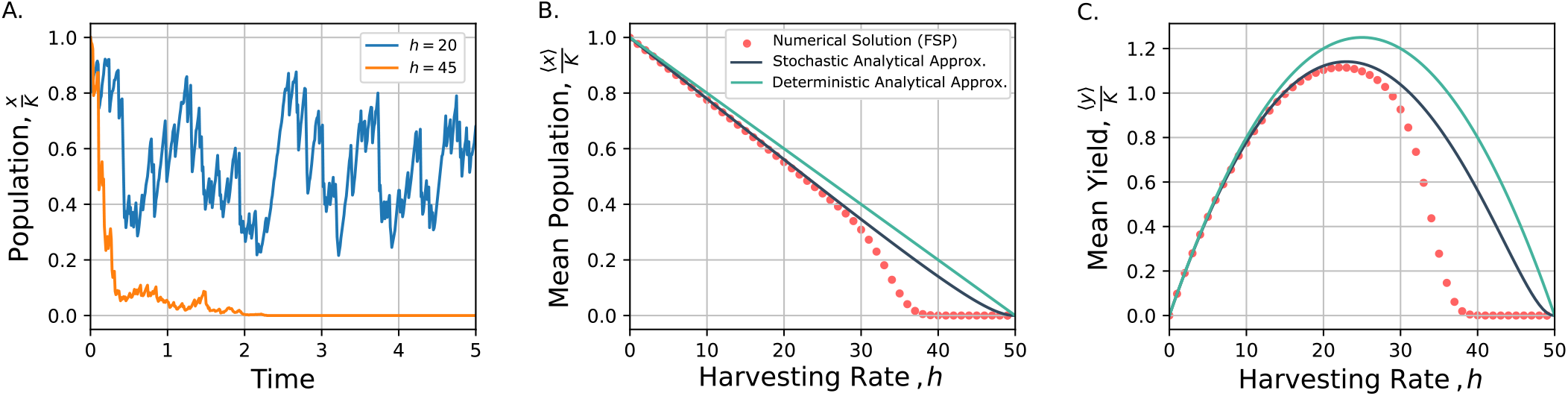
The total mean yield is maximized at an optimal harvesting rate. A. Two examples of normalized population trajectories over time are shown. Increasing the harvesting rate *h* leads the population to extinct. B. Increasing *h* for fixed *f* leads to the population extinction in a relatively shorter time. C. The maximum total mean yield is achieved at the optimal harvesting rate for fixed harvesting ratio *f*. The deterministic analytical result obtained from the ODE solution for 〈*x*〉 in (4). Additionally, the stochastic analytical result (blue) represents the outcome derived from the stochastic model proposed in section III-B. Red dots are collected from numerically computing the mean population and yield using finite state projection (FSP) method by solving (17) at a time *t_max_* = 400/*r*. Also, The values of 〈*x*〉 and 〈*y*〉 are normalized over the carrying capacity *K*. The rest of the parameters are taken as *r* = 10, *K* = 10^3^, *f* = 0.2, *λ* = 2.

Furthermore, in Fig. 4 we plotted the probability of extinction as a function of the harvesting rate *k* at a time *t* = 400/*r*, which is consistent with the results shown in Fig. 3. For the parameters considered in Fig. 3, the difference between the numerical results and the estimations from (31) is mainly due to a relatively high probability of extinction. Having a constant Poisson harvesting rate, the probability of harvest taking place is independent of the population count *x*. This lack of dependence on population count could contribute to a relatively early extinction. In the future work, we will propose the stochastic harvesting model with population-dependent harvesting rate to study how this strategy affects the extinction probability.

**Fig. 4:**
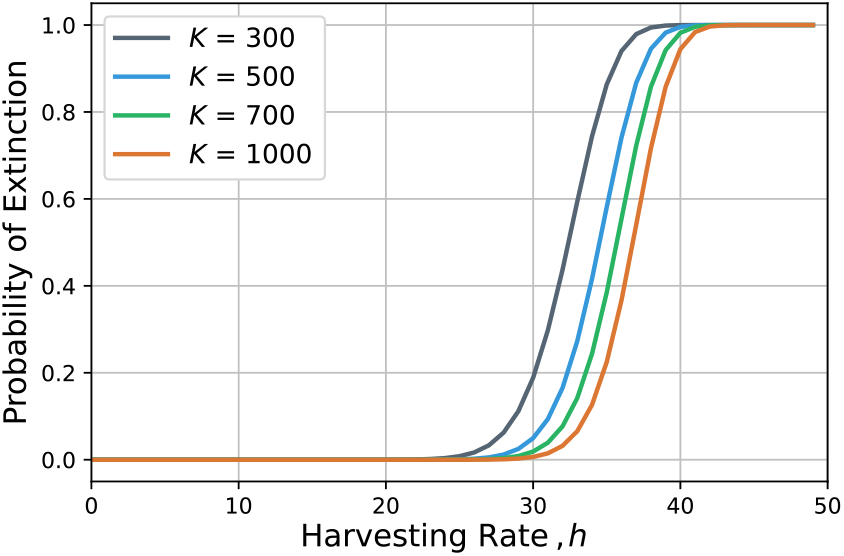
Probability of extinction increases as a function of the harvesting rate *h*. The probability of extinction approaches certainty (probability of 1) as the value of *h* increases, such that extinction becomes unavoidable. Curves are plotted for different *K* ∈ {300, 500, 700, 1000}, where *K* represents carrying capacity. In the simulation, we considered the time *t* in FSP as *t_max_* = 400/*r*. Other parameters are taken as *r* = 10, *f* = 0.2.

## IV. Conclusion

In this contribution we have used the framework of the stochastic discrete logistic model to study a specific class of environmental noise, where a fraction *f* of the population dies upon random occurrence of perturbation events. Exploiting the previously developed derivative-matching closure scheme, we developed analytical results quantifying the magnitude of fluctuations (11) that increases with *f* (after controlling for the deterministic steady-state mean). This implies that rare but more severe perturbation (as compared to frequent but less severe perturbations) can lead to larger random fluctuations, and these come at risk of higher extinction probability in a finite time. Then, we directly calculated this probability using the finite state projection (FSP) method, and numerically computed the probability of extinction as a function of *f* and perturbation frequency.

To investigate the harvesting problem, we generalized the model by including a yield variable *y* that is to be maximized, and we calculated the optimal harvesting rate. we found that a constant Poisson harvesting rate leads to extinction as the harvesting rate increases, but this did not occur at the optimal harvesting rate for the parameters used. Future work will include generalizing the model to consider a population-dependent harvesting rate and investigating the optimal harvesting strategy, which leads to the maximum yield.

## Notes

### Competing Interest Statement

The authors have declared no competing interest.

